# Tumor-myeloid crosstalk drives therapy resistance in localized bladder cancer

**DOI:** 10.1101/2025.09.08.674862

**Authors:** Filipe LF. Carvalho, Jihyun Lee, Nikolaos Kalavros, Yuzhen Zhou, Daniel Michaud, Isabella Stelter, Hiba Siddiqui, Konrad Stawiski, Ramya Ravindranathan, Julia Perera, Shuoshuo Wang, Breanna Titchen, Amanda Garza, Kevin Bi, Jihye Park, Elinor G. Sterner, Jillian M. Egan, Yuna Hirohashi, Raie T. Bekele, Ilana Epstein, Ioannis Vlachos, Li Jia, Adam S. Kibel, Michelle Hirsch, Joaquim Bellmunt, Jennifer L. Guerriero, Kent W. Mouw, Eliezer M. Van Allen

## Abstract

Neoadjuvant cisplatin-based chemotherapy results in pathologic complete response for only a minority of patients with muscle-invasive bladder cancer (MIBC), and mechanisms of resistance and the effects of chemotherapy on the MIBC microenvironment remain incompletely understood. Here, we defined the single-cell and spatial transcriptomes of cancer and immune cells from MIBC patients with resistance to cisplatin-based chemotherapy. Tumors with persistent MIBC after chemotherapy harbored cancer cells expressing epithelial-to-mesenchymal programs that were associated with worse overall survival in independent cisplatin-treated bladder cancer cohorts. These cisplatin-resistant tumor cells were infiltrated by macrophages that upregulated tumor permissive programs defined by increased PARP14 expression in spatially resolved multicellular niches. Macrophage reprogramming through PARP14 inhibition sensitized tumors to cisplatin via downregulation of tumor cell pathways implicated in resistance. Our results demonstrate that cancer cells and macrophages cooperate to promote cisplatin resistance and identify macrophage-directed PARP14 inhibition as a novel therapeutic strategy to sensitize MIBC to cisplatin.

## Introduction

Muscle-invasive bladder cancer (MIBC) is a potentially lethal disease state that often involves multimodal treatment with chemotherapy, surgery, and immunotherapy^1,2^. For many years, neoadjuvant cisplatin-based chemotherapy followed by radical cystectomy has been a standard curative-intent treatment for localized MIBC^3–5^. However, only 30% of MIBC patients experience pathologic complete response following neoadjuvant chemotherapy, and patients with residual muscle-invasive tumors at the time of radical cystectomy are more likely to develop metastatic bladder cancer and die from the disease^3^. Recent evidence suggests the combination of cisplatin-based neoadjuvant chemotherapy and immune checkpoint inhibition improves outcomes in MIBC^6^, implicating the bladder tumor microenvironment in mechanisms of response and resistance to cisplatin-based treatment. However, chemoimmunotherapy strategies only improve clinical outcomes in a subset of patients with MIBC. Thus, there is an urgent need to dissect the impact of cisplatin on tumor cells and the microenvironment to inform novel therapeutic approaches.

Genomic and transcriptomic characterization of response to neoadjuvant therapies in MIBC has largely focused on bulk-sequenced samples comprised of complex mixtures of tumor and non-tumor cell types^7–11^. Bulk RNA profiling broadly identified several molecular subtypes that reflect tumor heterogeneity, but do not yet guide clinical management^12,13^. Single cell analysis of preclinical bladder cancer models and localized tumors provided insights on the diversity of urothelial and non-urothelial cell types implicated in bladder cancer pathogenesis^14–18^. However, our understanding of how cytotoxic chemotherapy impacts bladder tumor cells and those in the microenvironment – particularly macrophages, which are the most abundant type of immune cells and contribute to tumor growth and immune evasion in this disease^19,20^ – remains limited in MIBC.

We hypothesized that specific interactions between malignant urothelial cells and immune cells jointly promote chemotherapy resistance in MIBC. To investigate this hypothesis, we paired complementary single-nucleus RNA sequencing (snRNAseq) and spatial transcriptomic methods in clinically integrated contexts with focused functional assessments to understand cancer and immune cell multicellular communities implicated in resistance to neoadjuvant cisplatin-based chemotherapy.

## Results

### Dissecting the cellular landscape of post-cisplatin bladder biopsies

We performed single-nucleus RNA sequencing (snRNA-seq) of frozen surgical resection specimens from MIBC patients treated with neoadjuvant chemotherapy followed by radical cystectomy (Fig. 1A, Suppl. Table 1). All patients received at least three cycles of cisplatin-based chemotherapy, and no patients had received prior intravesical therapy (e.g. BCG or intravesical chemotherapy), systemic anti-cancer therapy, or radiotherapy. All tumors were histologically assessed and reviewed by a board-certified genitourinary pathologist to confirm pure urothelial carcinoma without variant histology. Specimens were defined as “tumor scar” if pathology review indicated pathologic complete response (i.e., ypT0 N0) per established criteria^21^, or “persistent tumor” if there was persistent muscle-invasive (≥ypT2) disease and/or clinical progression to metastatic disease (Fig. 1B; Methods).

**Figure 1.**
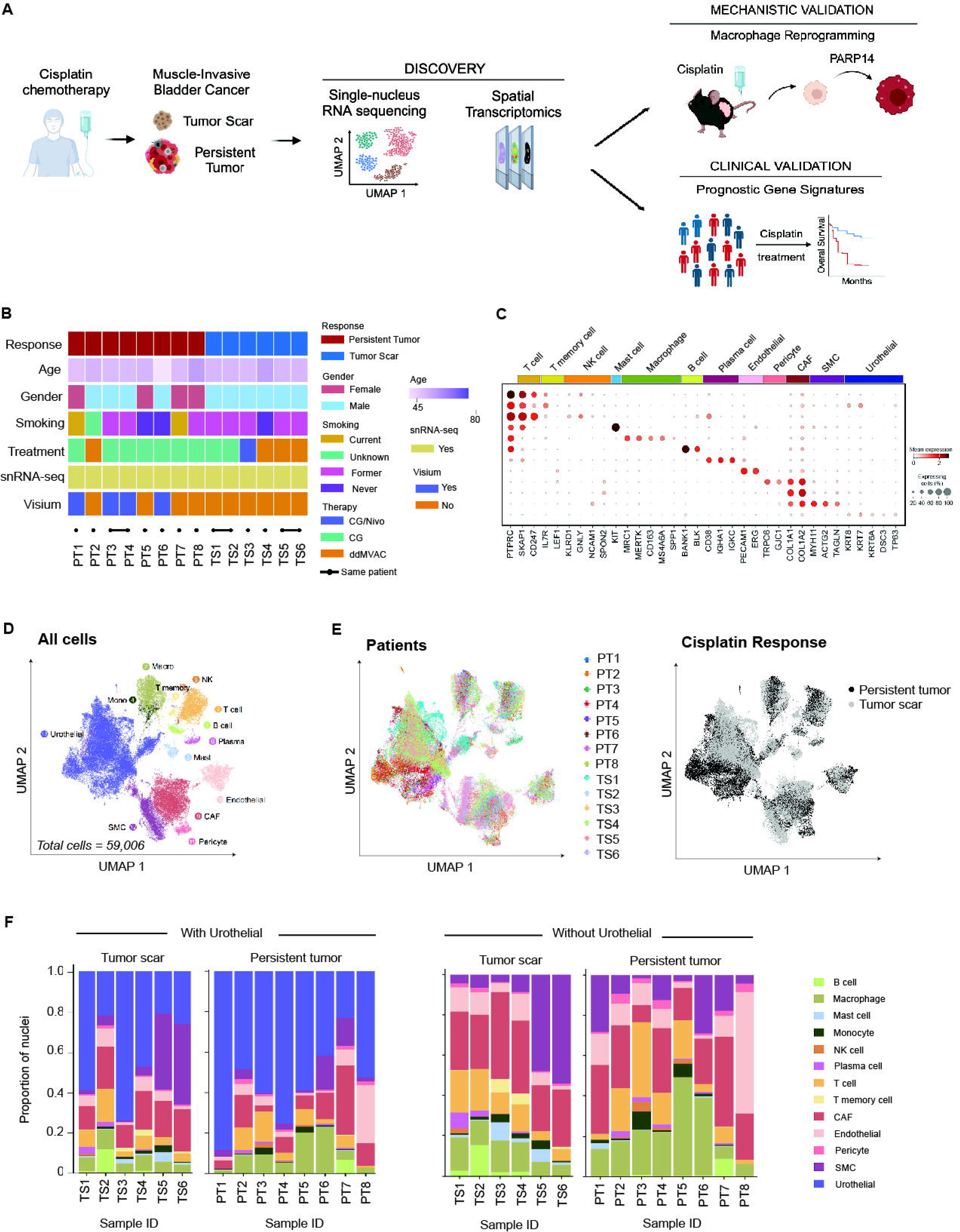
Defining the tumor microenvironment of cisplatin-treated MIBC. (A) Study overview and experimental workflow. Figure created with BioRender.com. (B) Summary of clinicopathological characteristics, treatment history and molecular single-nucleus and spatial transcriptomic profiling (Visium, 10x Genomics) of tumor samples. Treatment groups: PT, persistent tumor; TS, tumor scar. Treatment history: CG, cisplatin/gemcitabine; ddMVAC, Dose-Dense Methotrexate, Vinblastine, Doxorubicin, and Cisplatin; Nivo, Nivolumab. snRNA-seq, single-nucleus RNA-sequencing. (C) Bubble plot of mean expression and percentage of cells expressing selected marker genes (columns) across urothelial and non-urothelial cells (rows) in all samples. NK, natural killer cell; CAF, cancer-associated fibroblast; SMC, smooth muscle cell. (D) Uniform Manifold Approximation and Projection (UMAP) of epithelial, immune, and stromal cell subpopulations captured across all samples colored by broad cell type. Mono, monocytes; Macro, macrophages; NK, natural killer cell; Plasma, plasma cell; SMC, smooth muscle cell; Mast, mast cell; CAF, cancer-associated fibroblast. (E) UMAP of malignant and non-malignant cells captured across all samples colored by patient and response to cisplatin. (F) Bar chart with the proportion of cell types across samples. Cell subsets across persistent tumors and tumor scar either with (left) or without (right) urothelial cells.

After quality control, our single-cell atlas included transcriptomes from 59,006 nuclei spanning malignant and non-malignant cell types. To minimize batch effects across patients, tissue collection, and processing, all nuclei were integrated using Harmony^22^ and jointly clustered in an unsupervised manner. Clusters were then annotated based on expression of established marker genes^14,23,24^(Fig. 1C, Suppl. Fig. S1A). We identified 13 cell subsets comprising urothelial cells as well as non-urothelial cells present in the tumor microenvironment that were shared among patients, treatment and response groups (Fig. 1D–E, Suppl. Fig. S1B). We confirmed malignant cells in the urothelial compartment of persistent tumors by inferred copy-number alterations (CNAs), which were consistent with genomic profiles from previously described chemotherapy resistant urothelial tumors^11^(Suppl. Fig. S1C, S1D). Other identified cell types included non-malignant urothelial cells, immune cells, and diverse stromal cell types (cancer-associated fibroblasts, endothelial cells, pericytes and smooth muscle cells), consistent with the histologic composition of MIBC. snRNA-seq captured epithelial, fibroblast and immune cell populations across individual tumors and response groups (Fig. 1F). Within the immune cell compartment, we identified lymphoid and myeloid cell fractions in tumor scar and persistent tumors with robust representation for further interrogation.

### Tumor-intrinsic programs are implicated in cisplatin resistance and clinical outcome of MIBC

Gene expression signatures derived from bulk RNA profiling of MIBC tumors broadly divide tumors into luminal and basal subtypes, but the implications of these (or other) tumor-intrinsic transcriptional programs in predicting response to neoadjuvant therapies, resistance to cisplatin, and overall survival remain uncertain^13,25–27^. We hypothesized that tumor-intrinsic cellular programs enriched in persistent tumors may identify malignant cell features driving resistance to cisplatin. We first identified 30,413 putative urothelial cells based on criteria noted above, as well as cytokeratin (e.g. *KRT5*, *KRT6A*) and *TP63* expression across all profiled samples (Fig. 2A). We next used consensus nonnegative matrix factorization (cNMF^28^) to decompose expression profiles of single urothelial nuclei into programs present in the tumor scar and/or persistent tumors (Fig. 2B, Suppl Fig. S2A, Methods). We identified seven programs that reflect cell lineages (Myogenesis, EMT) or cell states (tumor necrosis factor/nuclear factor κB, phosphoinositide 3-kinase (PI3K)/ mammalian target of rapamycin (mTOR), interferon, drug metabolism, oxidative phosphorylation) (Fig. 2C, Suppl. Table 2). Most programs were detectable in both the tumor scar and persistent tumor cell populations (Suppl. Fig. S2B). Notably, the EMT program was significantly enriched in malignant urothelial cells from persistent tumor samples but largely inactive in the tumor scar (p = 0.039, two-sided Wilcoxon Rank Sum, Suppl. Fig. S2C). The cell surface marker CD44, previously implicated in bladder tumor invasion and metastasis^29^, was the most highly expressed gene in this EMT program. The EMT program was also enriched for genes implicated in cell proliferation and migration (e.g. *VOPP1*)^30,31^, as well as genes associated with MIBC basal/squamous differentiation (e.g., *CD44*, *TP63*)^12^, a cell phenotype induced by cisplatin in preclinical models of MIBC^32^ (Suppl. Fig. S2D).

**Figure 2.**
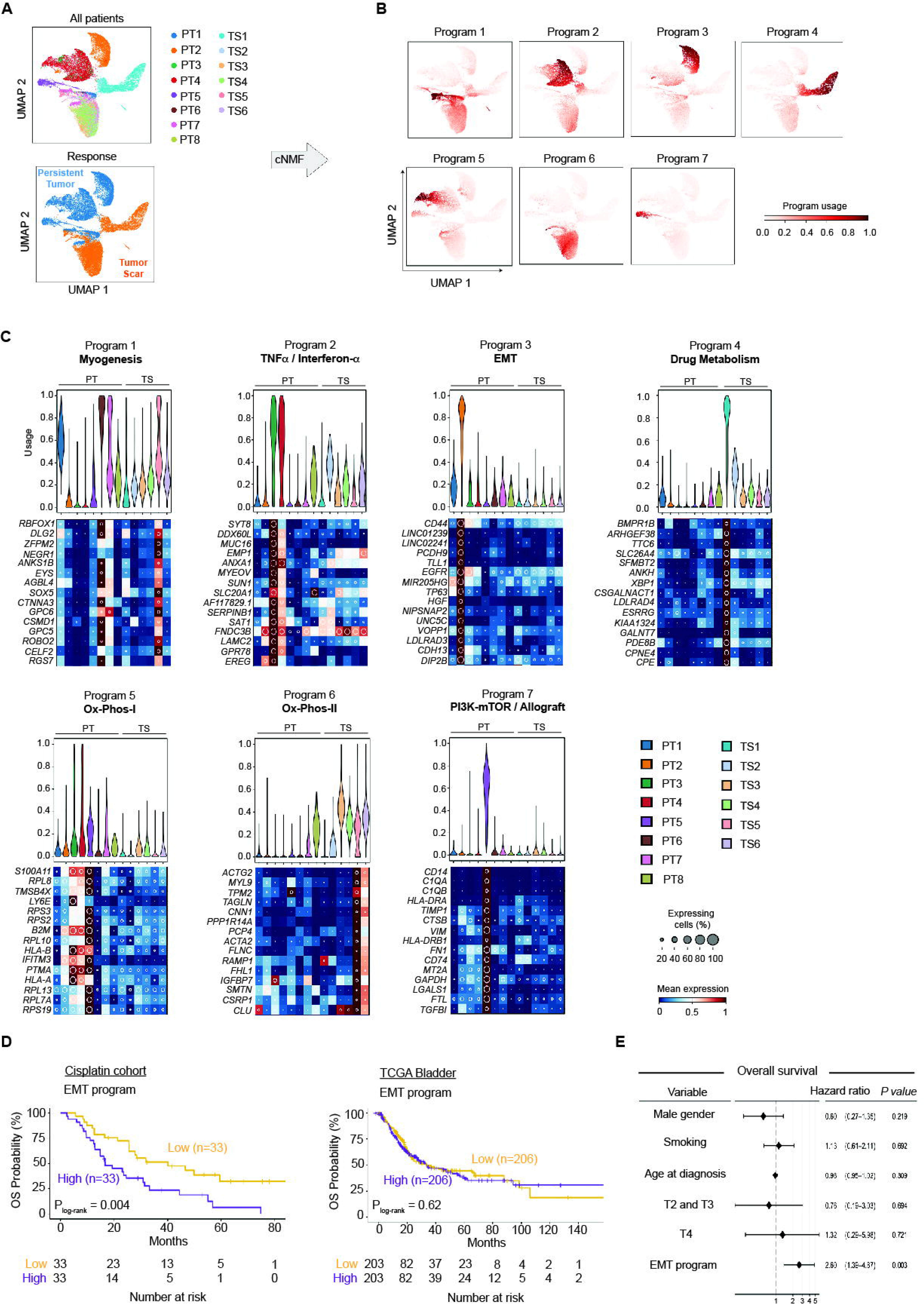
Malignant cell programs shared across persistent tumors are prognostic in cisplatin treated bladder cancer. (A) UMAP of urothelial cells captured across all samples colored by patient and response to cisplatin. (B) UMAP colored with per-cell relative usage of each of the seven expression programs identified by cNMF in urothelial cells. (C) Violin plots and heatmaps with distribution of cNFM usage program and expression level of the 15 top program-associated genes in each sample. (D) Kaplan-Meier analysis of overall survival (OS) in two independent cohorts: cisplatin-treated cohort (left) and TCGA treatment naïve MIBC patients (right). Patients were stratified by high and low EMT score in bulk RNA-seq. _Plog-rank_, log-rank test p value. (E) Multivariable Cox regression analysis of bulk RNA-seq data from cisplatin cohort incorporating gender, smoking status, age at diagnosis, tumor stage and EMT program expression as covariates. Data is presented as hazard ratio (HR) ± 95% confidence interval (CI).

Whereas the EMT program was preferentially enriched in the persistent tumor cells, an Ox-Phos program was active in both persistent tumor and tumor scar. Prior bulk RNA-seq studies showed that cisplatin induces an overall expression of the Ox-Phos pathway in bladder cancer cell lines^33^. However, analysis of the single cell data allowed us to identify distinct subsets of Ox-Phos genes expressed in malignant urothelial cells present in persistent tumors versus benign urothelial cells in the tumor scar (Ox-Phos-I and Ox-Phos-II; Fig 2C). Overall, these results suggest cisplatin induces common gene expression programs in both benign and malignant urothelial cells, while also driving distinct tumor-specific programs in malignant cells from non-responders that may reflect interactions with the tumor microenvironment.

To assess the potential predictive and prognostic implications of these malignant-cell-specific programs enriched in persistent tumor compared with tumor scar, we projected these programs into clinically annotated bulk RNA-seq data from two large, independent MIBC cohorts: (i) 66 patients with primary locally advanced or metastatic bladder cancer treated with cisplatin-based chemotherapy^34^ and (ii) 412 patients with primary MIBC not treated with neoadjuvant therapy from The Cancer Genome Atlas (TCGA)^12^. Bulk tumor samples from the cisplatin treated patients were profiled prior to initiation of systemic chemotherapy. We focused on the three programs (EMT, Ox-Phos-I usage in persistent tumors, and Ox-Phos-II usage in tumor scar) that were significantly different between the tumor scar and persistent tumors in our discovery cohort (p < 0.05, two-sided Wilcoxon Rank Sum, Suppl. Fig. S2C). We observed an association between pre-treatment EMT score and significantly worse overall survival (OS) in the cisplatin-treated cohort but not in the untreated (TCGA) cohort (P = 0.004, log-rank, Fig. 2D). No significant associations were found between Ox-Phos-I or II programs and survival in patients in either cohort. We then performed a multivariable Cox regression analysis of OS with age, sex, smoking, tumor stage, and the malignant EMT program as covariates. The EMT program (HR 2.60; 95% CI 1.39-4.87; p < 0.01) remained significantly associated with shorter OS in the cisplatin treated cohort (Fig. 2E, Suppl. Fig. S3A, S3B). These results support an association between an EMT gene signature and poor outcomes in MIBC patients treated with cisplatin-based chemotherapy. Altogether, these findings show that tumor-intrinsic urothelial programs from cisplatin-resistant cells exhibit distinct biological states that may have therapeutic implications.

### SPP1/PARP14 macrophage polarity defines the microenvironment of cisplatin resistant MIBC

We next examined whether the EMT-enriched persistent tumor cells harbored unique immune cells in the tumor microenvironment. Prior studies have shown that cisplatin is associated with T cell priming, and the combination of cisplatin-based chemotherapy with anti-PD-L1 immune checkpoint blockade has been shown to improve OS in MIBC and metastatic bladder cancer compared to chemotherapy alone^6,35^. Thus, we compared the distributions of immune cells in tumor scar and persistent tumor, noting that T cells and tumor-associated macrophages (TAMs) were the immune cell populations most represented in our cohort (Fig. 3A).

**Figure 3.**
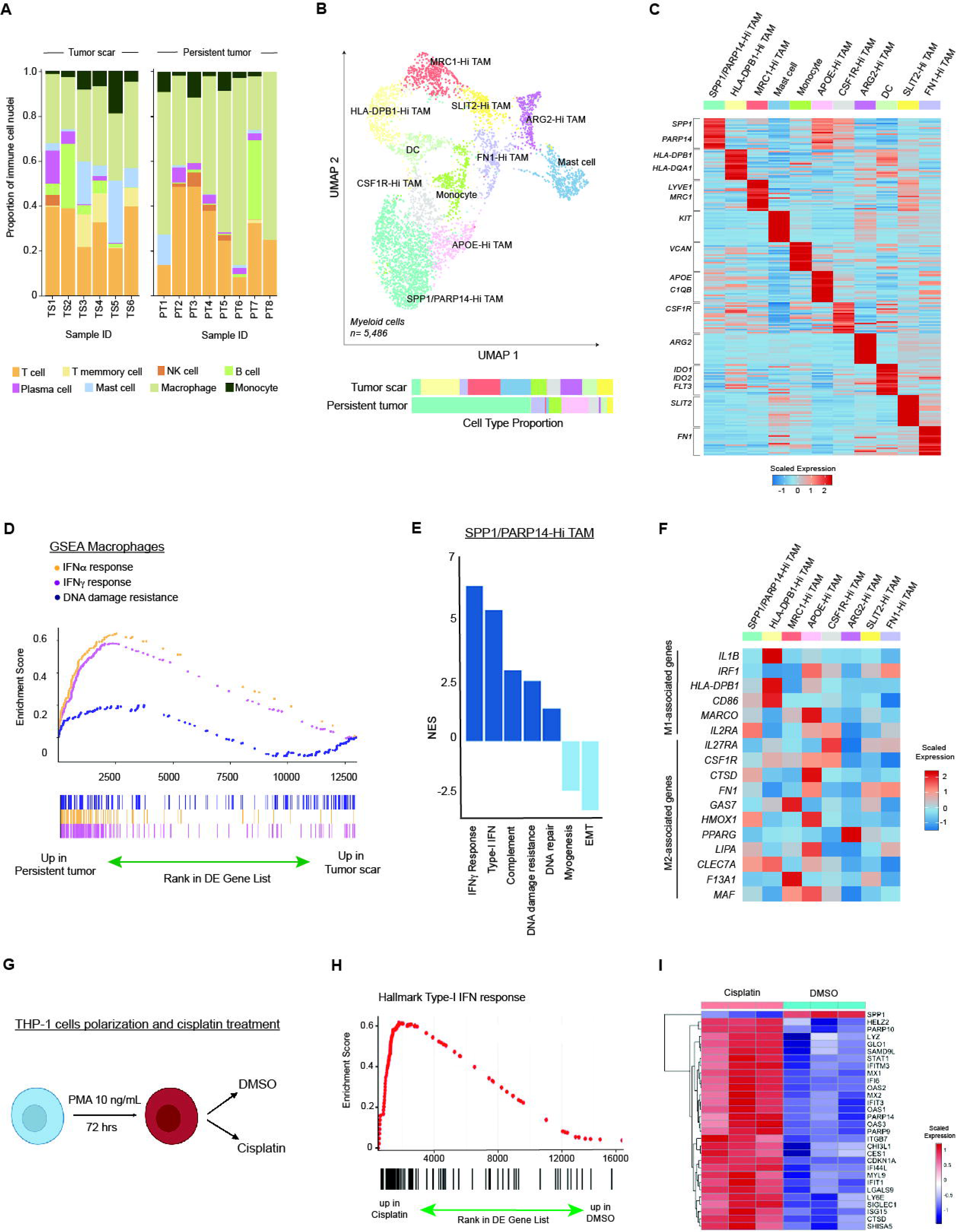
SPP1/PARP14 TAMs characterized by interferon activation and DNA damage resistance signatures are enriched in cisplatin-resistant tumors. (A) Bar chart with the proportion of immune cells across persistent tumors and tumor scar. (B) UMAP of myeloid cells captured across all samples, labeled and colored by cell type. Bar plots show cell type proportions grouped by response to treatment. (C) Heatmap of scaled normalized expression for cell type defining genes determined by two-sided Wilcoxon rank-sum test with Bonferroni FDR correction (q < 0.05). (D) GSEA of Hallmark interferon-γ, interferon-α and a validated interferon-related gene signature for DNA damage resistance^48^ in macrophages from persistent tumor and tumor scar. (E) Bar chart with normalized enrichment scores (NES) of Hallmark and a literature curated gene set^48^ significantly enriched and depleted in SPP1/PARP14 TAMs. Two-sided Wilcoxon rank-sum test with Bonferroni FDR correction, q < 0.05. (F) Heatmap of scaled normalized expression of curated genes within macrophage subpopulations. (G) Overview of the experimental workflow of THP-1 cells polarized and treated with cisplatin *in vitro*. Figure created with BioRender.com (H) GSEA of Hallmark interferon-α in polarized THP-1 cells treated with DMSO and cisplatin. (I) Heatmap of scaled normalized expression most highly differentially expressed genes in THP-1 cells treated with cisplatin compared with DMSO determined by two-sided Wilcoxon rank-sum test with Benjamini-Hochberg (BH) FDR correction (q < 0.05).

Myeloid cells, specifically TAMs, play an important role in the tumor microenvironment and contribute to treatment resistance in several tumor types^24,36–39^; however, their role in MIBC and cisplatin resistance remains uncertain. TAMs were the most abundant immune cell population across the cohort, and the overall proportion of TAMs in tumor scar compared to persistent tumor was not significantly different (Suppl. Fig. S4A). However, distinct subsets of TAMs are known to exhibit different transcriptional cell states and phenotypes, so we hypothesized that certain TAM subtypes drive selective cisplatin response in MIBC. Using differential gene expression and examination of known macrophage markers, we identified eight TAM clusters expressing distinct markers implicated in anti-tumor and pro-tumor roles in the microenvironment (Fig. 3B)^40^. The only TAM cluster significantly enriched in persistent tumors (p = 0.017, two-tailed t-test, Suppl. Fig. S4B) exhibited higher expression of the secreted factor SPP1 as well as PARP14, which has been implicated in macrophage polarization by directing an anti-tumor to pro-tumor phenotype shift (Suppl. Fig. S4C)^41,42^. The other seven macrophage clusters expressed genes that denote macrophage subpopulations with previously defined roles in inflammation (e.g. *HLA-DBP1, ARG2*)^43^, tissue metabolism (e.g. *APOE*) and fibrosis (e.g. FN1)^44^, as well as those present in the tumor microenvironment in other cancer types (e.g. *MRC1, SLIT2, FN1* and *CSF1R*)^23,37,45^ (Fig. 3C, Suppl. Table 3).

To complement TAM clustering analyses, we performed gene set enrichment analysis on the total TAM population to independently nominate programs enriched in the cisplatin resistant setting. TAMs in persistent tumors demonstrated upregulation of gene sets associated with interferon-γ (IFN-γ), which promotes macrophage activation; interferon-α (IFNα), which has a known role in the regulation of DNA damage response and acquired resistance to cisplatin^46,47^; and an interferon-related DNA damage resistance signature^48^ (Fig. 3D). Because SPP1/PARP14-Hi TAMs were the TAM subpopulation most enriched in persistent tumors, we hypothesized that this TAM subtype had a phenotype implicated in resistance to cisplatin. Indeed, we found that SPP1/PARP14-Hi TAMs were significantly enriched for interferon-α, hallmark DNA damage repair pathway, and DNA damage resistance^48^ signatures compared to other TAM subpopulations in persistent tumors (Fig. 3E; Methods).

Tissue macrophages adopt distinct phenotypes in response to stimuli that have historically been described as M1 pro-inflammatory or M2 tumor-permissive polarization states^49^. However, there is increasing recognition that macrophages undergo differentiation states along a continuum of phenotypes not fully captured by a strict dichotomization as “anti-tumor” M1 or “pro-tumor” M2^50^. HLA-DPB1-Hi macrophages that predominated in the tumor scar strongly expressed genes associated with an anti-tumor phenotype (*IL1B, HLA-DPB1, CD86*), whereas the SPP1/PARP14-Hi population that was enriched in persistent tumor predominantly expressed M2-associated genes such as *CTSD*, which has been implicated in promoting invasion and metastasis in breast cancer^51^ (Fig. 3F).

To experimentally validate our findings that macrophages in cisplatin resistant tumors activate type-I IFN signaling, we polarized human THP-1 cells and treated with cisplatin (Fig. 3G; Methods). Transcriptomic analyses revealed significant upregulation of type-I IFN signaling in THP-1 cells treated with cisplatin (Fig. 3H), and the pattern of cisplatin-induced upregulation of these type-I interferon-response genes (ISG) (e.g. *IFI6, OAS2, OAS3, IFIT1, IFIT3, IFITM3, ISG15*), as well as *STAT1 and PARP14* (Benjamini-Hochberg adj p < 0.05), matched the expression profiles of the SPP1/PARP14-Hi TAM population in persistent tumors (Fig. 3I). Notably, treatment with cisplatin did not increase SPP1 expression in THP-1 cells, a result consistent with prior observations in other cancer types (e.g. colorectal cancer) where macrophages cultured *in vitro* are unable to express SPP1 because direct cell-cell contact between tumor epithelial cells and macrophages is required to induce SPP1 expression^52^. Overall, our results relate resistance-associated tumor-intrinsic programs, such as EMT, to the surrounding microenvironment characterized by activated SPP1/PARP14-Hi TAMs, that together may contribute to cisplatin resistance.

### Cancer cell-TAM interactions are spatially organized in MIBC

Given our observation that *CD44* was the most highly expressed gene in the malignant cell EMT program that was associated with worse survival in cisplatin-treated bladder cancer cohorts, as well as our observation of *SPP1* upregulation in TAMs in persistent tumors, we hypothesized that CD44-expressing cancer cells and SPP1-expressing macrophages cooperate to promote cisplatin resistance. We first analyzed our snRNAseq cohort using CellphoneDB^53^ to identify candidate cellular crosstalk between urothelial cells and all macrophage subpopulations through known ligand-receptor pairs (Suppl. Fig. S5). Non-cytotoxic macrophages (e.g, HLA-DBP1-Hi) expressed *IL-1* and TNF programs that have been implicated in the inflammatory response as well as metalloproteinase genes that may maintain tissue homeostasis in both the tumor scar and persistent tumors. However, only malignant urothelial cells enriched in the EMT program in persistent tumors were inferred to use CD44 to engage SPP1 on TAMs, an interaction previously implicated in initiation of bladder cancer metastasis in mouse models^29^.

Based on these observations, we hypothesized that urothelial cells and SPP1/PARP14-Hi macrophages define spatially organized multicellular communities in persistent tumors. To further investigate co-localization of macrophages with tumor cells and interrogate cellular crosstalk, we performed *in situ* whole transcriptome analysis of a subset of persistent tumors that also underwent single cell analysis (Methods). The expression of epithelial, stromal, and immune cells appropriately mapped the histologic location of these cell types as assessed by genitourinary pathologist review (Fig. 4A, Suppl. Fig. S6, Suppl. Table 4, and Methods). At the cell subtype level, we found an enrichment of urothelial cells in the epithelial compartment and a predominance of smooth muscle cells in the stromal compartment, consistent with the anatomical location and biology of MIBC (Fig. 4B; Methods). Moreover, TAMs were the predominant immune cell population present in the tumor microenvironment (Fig. 4B). We therefore applied the snRNA-seq SPP1/PARP14-Hi-specific signatures in spatial gene-expression based clustering to define the location of this population of TAMs in bladder tumors. We confirmed that SPP1/PARP14-Hi TAMs were significantly enriched in two spatially defined clusters (IMM_1 and IMM_2, adjusted p < 0.001, two-sided Wilcoxon Rank Sum Test with Holm-Bonferroni FDR correction, Fig. 4C).

**Figure 4.**
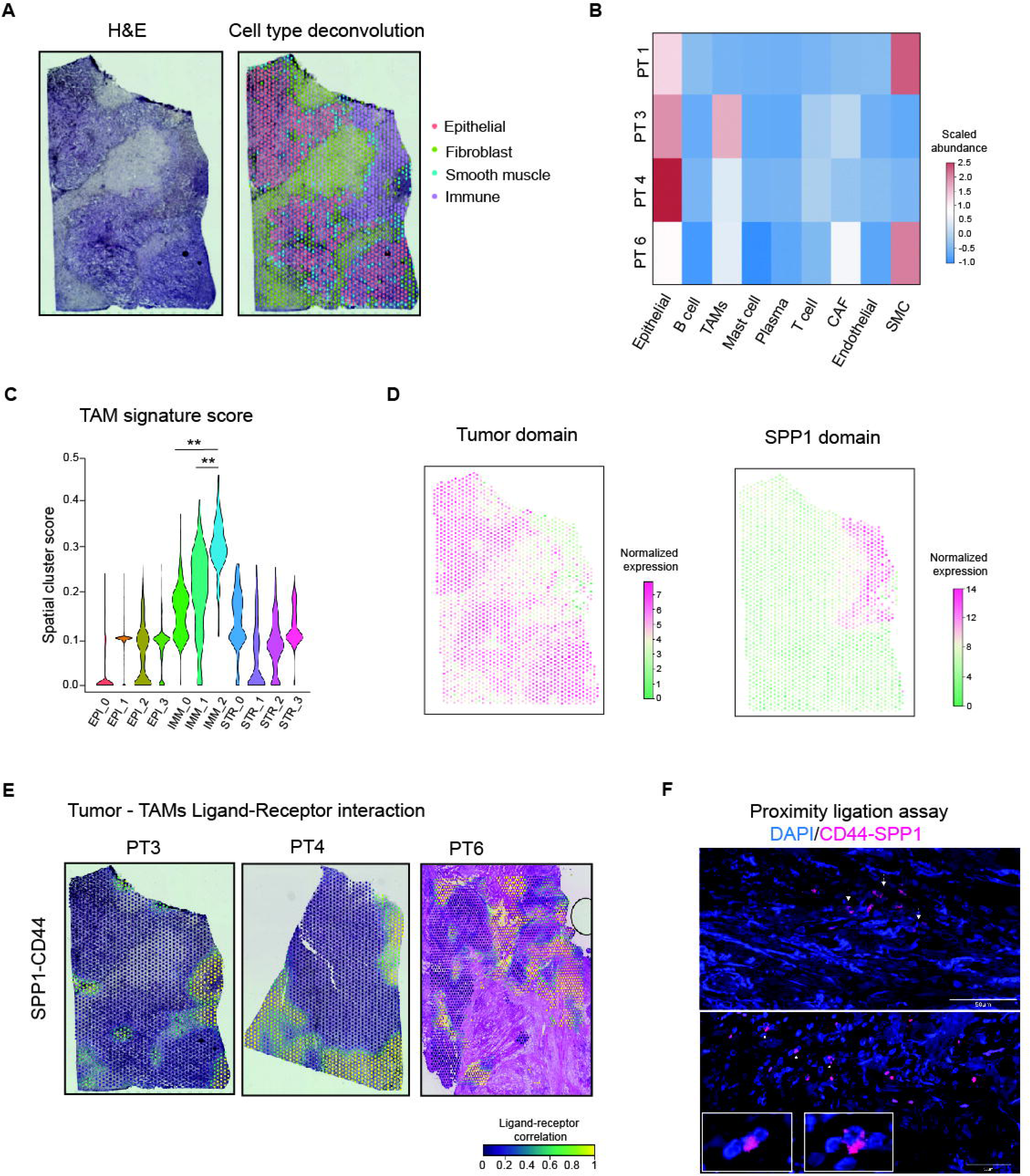
Malignant urothelial cells and SPP1 TAMs define spatially restricted cellular neighborhoods in cisplatin resistant tumors. (A) Representative hematoxylin and eosin (H&E) and coarse spatial mapping of cell types per spot in the same bladder tumor using established lineage markers (as in Fig. 1C). (B) Heatmap with cell type abundance across samples. Granular cell types were discerned in each spot using patient-matched single-nucleus references and cell-type specific gene markers. TAMs, Tumor-associated macrophages; CAF, cancer-associated fibroblasts; SMC, smooth muscle cells. (C) Violin plot with TAMs single-nucleus gene signatures across spot-defined cell populations (**adjusted p < 0.001, two-sided Wilcoxon Rank Sum Test with Holm-Bonferroni FDR correction). (D) Spatial domains with normalized expression of SpaGCN inferred signature defined tumor and SPP1-high TAM regions (pink spots). (E) Cell-cell interaction maps of CD44-SPP1 in cancer cells and TAMs identified using LIANA+. Single spots with higher co-expressed CD44-SPP1 interaction (yellow) in spatial contexts in representative sections of persistent tumors. (F) Representative immunofluorescence images of the CD44–SPP1 proximity ligation assay (PLA) detected in two distinct persistent tumors. CD44–SPP1 protein–protein interactions are visualized as fluorescent signal (pink) in adjacent cells, consistent with ligand–receptor engagement. The bottom panel shows a magnified view of the region highlighted with arrows in the upper image.

To further examine the spatial co-existence between SPP1/PARP14-Hi TAM and persistent tumor cells, we used SpaGCN^54^ to determine the proximity between these two cell types in spatially restricted tumor areas. We first identified tumor spatial domains defined by *KRT8* expression overlaying histological areas of tumors cells (Fig. 4D, left panel). In each case, this tumor domain was adjacent to a neighboring TAM spatial domain defined by the expression of *SPP1*, *S100A8*, which drives monocyte differentiation towards pro-tumor TAMs, and *IFI6*, a type-I IFN-stimulated gene expressed in the pro-metastatic tumor microenvironment^46^ (Figs. 4D, 3I).

We then further dissected tumor-TAM crosstalk in macrophage-enriched spatial domains by defining the correlation between expression of ligand genes in macrophages and receptor genes in neighboring cancer cells^55^. This cell-cell communication analysis in tumors sections with high macrophage content identified preferential CD44-SPP1 ligand-receptor pairs in macrophage enriched spatial domains (Fig. 4E). This predicted CD44-SPP1 interaction in persistent tumors was validated using multiplex immunofluorescence and a proximity-ligation assay to visualize interactions between CD44-expressing tumor cells and SPP1-expressing TAMs (Fig. 4F; Suppl. Fig S7A – S7B; Suppl. Table 5; Methods). These results identify multicellular niches of cancer cells and macrophages characterized by the crosstalk between SPP1/PARP14-Hi TAMs and CD44-expressing tumor cells in MIBC resistant to cisplatin.

### Macrophage reprogramming through PARP14 inhibition

Macrophages can be therapeutically targeted through: (i) activation blockade, using small molecule inhibitors and antibodies to inhibit their recruitment to the tumor; (ii) depletion of tumor promoting TAMs; and (iii) reprogramming of tumor promoting TAMs to an anti-tumor phenotype^38,40^. Given our finding that SPP1/PARP14-Hi TAMs co-localize with cancer cells in cisplatin-resistant bladder tumors and because PARP14 is implicated in pro-tumor macrophage polarization^41,42^, we hypothesized that inhibition of PARP14 could reverse this tumor permissive TAM phenotype and potentiate the anti-tumor effect of cisplatin.

We first assessed the impact of cisplatin exposure on PARP14 expression in murine RAW264.7 macrophages and observed a dose-dependent increase in PARP14 expression (Fig. 5A), consistent with results observed in human THP-1 cells (Fig. 3I). RBN012759 is a selective inhibitor of PARP14 (PARP14i) that has been shown to reverse pro-tumor macrophage gene expression programs and restore sensitivity to immune checkpoint blockade in kidney cancer and melanoma^42,56^. Treatment of RAW264.7 macrophages with RBN012759 caused a significant increase in PARP14 expression, as has been previously described^42^, and this effect was further increased by co-treatment with IFNγ, which has also been demonstrated to increase PARP14 expression^56^ (Fig. 5B, Suppl. Fig. S8A). To validate the activity of RBN012759 in suppressing PARP14 signaling, we performed immunoblotting and observed an increase in phospho-STAT1 expression in RAW264.7 macrophages treated with RBN012759, consistent with relief of PARP14-mediated STAT1 suppression^56^, as well as inhibition of IL4-driven expression of the pro-tumor macrophage markers Cd206 and Arg1 (Fig. 5C). Similar effects were observed in human THP-1 macrophages treated with cisplatin plus PARP14i *in vitro* (Suppl. Fig. S8B). Together these data demonstrate that cisplatin directly upregulates macrophage PARP14 expression and that PARP14 inhibition with RBN012759 inhibits PARP14-mediated signaling in macrophages.

**Figure 5.**
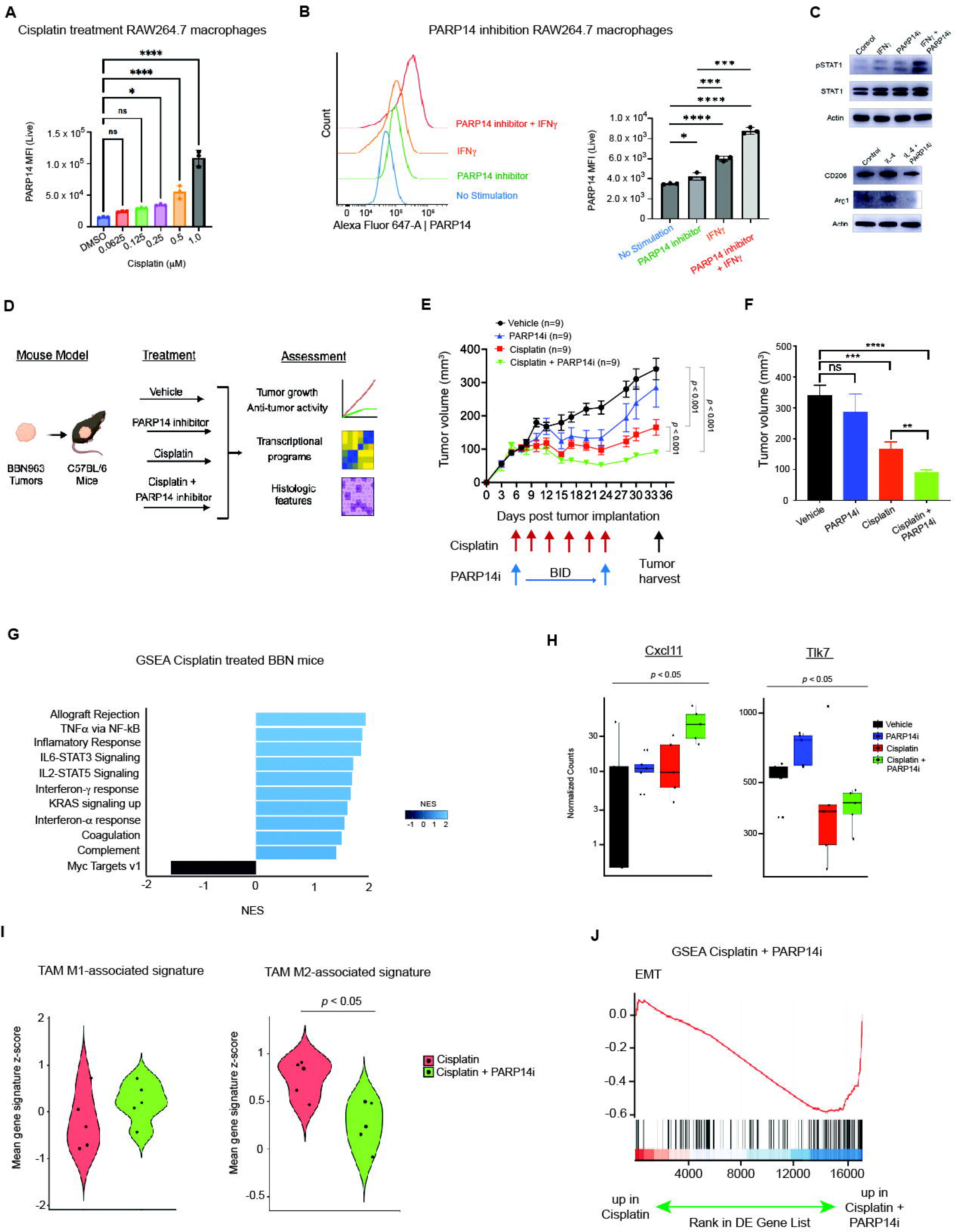
PARP14 Inhibition combined with cisplatin reduces tumor growth in preclinical models of bladder cancer through downregulation of EMT programs. (A) Bar chart with PARP14 expression in RAW264.7 mouse macrophages with increasing concentrations of cisplatin measured by flow cytometry (one-way ANOVA with Benjamini-Hochberg (FDR) correction (*q < 0.05; ****q < 0.0001)). (B) Histogram (left) and bar chart (right) of PARP14 expression in RAW264.7 macrophages treated with DMSO, PARP14i, IFN-γ or PARP14i + IFN-γ (unpaired two-tailed t-test (*p < 0.05; ***p < 0.001; ****p < 0.0001)). (C) RAW264.7 mouse macrophages treated with IFNγ, IL-4 and PARP14 inhibitor for 24 hours. The protein expression of the PARP14 regulated genes pSTAT1, STAT1, CD206 and Arg1 was determined by western blot with β-actin (*ACTB*) used as loading control. (D) Experimental design, treatment groups and measures of mouse xenografts. (E) Tumor volume data for mice treated with vehicle, PARP14i, cisplatin or cisplatin + PARP14i. (F) End-of experiment (day 34) tumor volumes for mice treated with vehicle, PARP14i, cisplatin, or cisplatin + PARP14i (unpaired two-tailed t test, **p < 0.01; ***p < 0.001; ****p < 0.0001). (G) Bar chart with normalized enrichment scores (NES) of MSigDB^62^ gene sets significantly enriched and depleted in tumors from mice treated with cisplatin. Two-sided Wilcoxon rank-sum test with Bonferroni FDR correction, q < 0.05. (H) Box plot with normalized counts for the PARP14 target genes Cxcl11 and Tlk7 in mouse tumors across treatment groups (Krushkal-Wallis test, p < 0.05). (I) Violin plots with normalized expression of published M1- and M2-associated gene signatures^58^ in tumors treated with cisplatin or cisplatin + PARP14i. (J) GSEA of Hallmark EMT gene set in tumors treated with cisplatin or cisplatin + PARP14i.

We next investigated the effects of cisplatin and PARP14i treatment on tumor growth and microenvironmental changes in a syngeneic immune-competent mouse model of bladder cancer (Fig. 5D). BBN963^57^ tumor-bearing mice were randomized to treatment with vehicle, cisplatin, PARP14i, or cisplatin plus PARP14i. Treatments were well tolerated, with no significant changes in mouse body weights across treatment arms (Suppl. Fig. S9A). Cisplatin monotherapy resulted in a significant decrease in tumor size compared with vehicle control (unpaired two-tailed t test, p < 0.001), whereas PARP14i monotherapy did not impact tumor growth. Mice treated with cisplatin plus PARP14i demonstrated a significant reduction in tumor volume compared to cisplatin monotherapy (unpaired two-tailed t test, p < 0.01, Fig. 5E – F).

At the end of the experiment, mice were sacrificed, and bulk RNA sequencing of tumors was performed (Methods, Suppl. Fig. S9B – S9C) to assess expression of genes known to be modulated by RBN012759. Cisplatin monotherapy led to increased expression of multiple inflammatory gene sets compared to vehicle (TNFα, type I IFN and IFN-γ) (Fig. 5G), similar to the gene sets upregulated in TAMs and urothelial cancer cells from persistent tumors from MIBC patients (Fig. 2C, 3D-E). Moreover, chemokines such as *CXCL11* and *TLR7* that promote an anti-tumor microenvironment and are regulated by PARP14 in preclinical models of kidney cancer^42^ and in human THP-1 cells (Suppl. Fig. S8B) were also significantly upregulated in mouse tumors treated with cisplatin plus PARP14i compared with cisplatin alone (Fig. 5H).

We next used the RNA sequencing data to assess established macrophage gene signatures implicated in tumor-permissive (M2-associated genes) and anti-tumor (M1-associated genes) phenotypes^58^. Although the total number of macrophages was similar across treatment arms as assessed by flow cytometry (Suppl. Fig. S9D), the addition of PARP14i to cisplatin resulted in a significant downregulation of an M2-associated gene signature (two-sided Wilcoxon rank sum test, p < 0.05) as well as increase in an M1-associated signature (Fig. 5I), consistent with a shift in macrophage polarization from a tumor-permissive to an anti-tumor phenotype. In addition, the combination of cisplatin with PARP14 inhibitor caused a significant downregulation of EMT gene programs (Fig. 5J) that were high in the persistent tumors from MIBC patients (Fig. 2C-E). Finally, complement and myogenesis pathways were also downregulated in cisplatin plus PARP14i-treated tumors, consistent with suppression of PARP14 leading to anti-tumor effects in both cancer and immune cells^41,42,56^ (Suppl. Table 6). Collectively, these results demonstrate that the combination of cisplatin plus the PARP14i RBN012759 significantly decreases tumor growth and that PARP14 inhibition elicits anti-tumor immune gene expression to promote an inflammatory anti-tumor microenvironment in MIBC.

## Discussion

To define cancer and immune cell interactions responsible for cisplatin resistance, we comprehensively analyzed single nuclei and spatial transcriptomes from bladders of patients with and without persistent MIBC after chemotherapy. We uncovered populations of chemotherapy-refractory cancer cells defined by the expression of multiple transcriptional programs. Tumor cell intrinsic EMT gene signatures derived from our single cell dataset were associated with worse overall survival when applied to bulk RNA sequencing data from two independent cohorts of cisplatin treated patients. These results suggest that a transition between epithelial and mesenchymal cell states contributes to therapy resistance and that EMT signatures may identify patients most likely to benefit from cisplatin-based chemotherapy. Within the myeloid cell compartment, a subpopulation of TAMs expressing *SPP1* and *PARP14* was enriched in persistent tumors. These TAMs expressed DNA damage resistance-related genes (e.g. *CTSL*^59^) as well as type-I interferon and cysteine cathepsins that have been previously associated with resistance to cytotoxic chemotherapy^48,60,61^. Furthermore, we observed that SPP1/PARP14-Hi TAMs co-existed in niches with cancer cells expressing EMT programs in persistent tumor samples. Analysis revealed significant cell-cell interactions within multicellular niches, where TAMs express ligands for interferon-γ induced cytokines (IL-18) implicated in macrophage activation as well as chemokines (CXCL16), integrins (ICAM, TIMP2), complement (C1QB) and secreted factors (SPP1) that foster cancer cell survival and resistance to therapy^38^. Macrophages treated with cisplatin exhibited a dose-dependent increase in PARP14 expression, and the addition of PARP14 inhibitor to cisplatin decreased tumor growth and downregulated signaling pathways (e.g. EMT) and reprogrammed TAMs to an anti-tumor phenotype in mouse bladder cancer models. Overall, these results support a model of macrophage reprogramming and establish PARP14 as a rationale target to overcome cisplatin resistance in MIBC.

There were several limitations to this study. Bladder cancer is a complex mixture of cancer cells, immune and stroma, and multi-regional sampling would be needed to fully capture tumor heterogeneity. In addition, we profiled frozen samples for snRNA-seq which present with technical challenges associated with cell degradation and single-nucleus extractions. Nonetheless, the cancer-immune cell interactions identified in persistent tumor samples were consistent across patients and anatomic tumor locations. Profiling matched pre- and post-cisplatin treated tumors can also inform the progressive tumor and immune cell phenotypes over cytotoxic chemotherapy. Nevertheless, we validated our findings in pre-treatment samples from bulk-sequencing cohorts, including one longitudinal prospective clinical trial, that confirmed the relevance of single-cell results. Additionally, pure urothelial carcinoma is the most common histologic type of urothelial cancer, so we purposely excluded variant histology tumors given the sparsity of data in cisplatin treated MIBC. Moreover, although we included both male and female patients, there was a predominance of female patients in the persistent tumor group, but all patients in the tumor scar group were male. Future work could focus on larger cohorts with balanced numbers of patients from both genders and include variant histology urothelial carcinoma to decipher convergent mechanisms of resistance to therapy. Lastly, our work focused on tumor-immune cell interactions, but the stromal compartment may also play an important role in resistance to therapy in MIBC that should be investigated in future studies.

Our results provide a roadmap for understanding cancer-immune cell interactions and modulation of macrophage phenotypes to sensitize MIBC to cytotoxic chemotherapy. Cisplatin based chemotherapy is a backbone of systemic therapies in MIBC, and the combination of cisplatin and anti-PD-L1 immunotherapy has recently been shown to improve outcomes in both localized MIBC and metastatic urothelial cancer^6^. This work identifies a distinct TAM population that contributes to cisplatin resistance and can be reprogrammed through PARP14 inhibition. This approach could potentially be integrated with anti-PD-1 therapy to further drive anti-tumor responses and improve clinical outcomes.

## Methods

### Ethics and human patient specimens

All patient samples were collected under the Dana-Farber Cancer Institute IRB protocol #02-021. All patients in this study were consented without compensation at the time of their clinic appointments by a member of the surgery staff and asked if they were interested in participating in an excess tissue biobank protocol involving the collection of biological samples and clinical data. For inclusion in this protocol, patients had been diagnosed with localized bladder cancer and went surgical resection. The study was explained in detail to all patients, all questions were answered and informed consent signed to participate. Patients were specifically consented to (i) collect tissue, blood and urine; (ii) link clinical information to collected tissues; (iii) permission for the study staff to contact the patient in the future for potential studies they might be appropriate to participate; (iv) use samples for the creation of cell lines and testing with available or under development sequencing techniques; (v) samples to be stored indefinitely until used. A copy of the consent was provided to the patient and the consent status recorded and kept updated in the database. Resected primary bladder tumors was examined by a board-certified pathologist to confirm neoplastic content and then excess tissue frozen in OCT and stored at –80°C for single nucleus RNA-sequencing. Matched FFPE and OCT blocks were used for Visium (10x Genomics) in 4 cases.

### Single nucleus isolation from frozen samples

The following Tween-20-Salt-Tris (TST) nuclei isolation protocol was adapted from a previously described technique^64^ and optimized by us for bladder tumors. All reagents were prepared and maintained on ice. A 2x stock ST was prepared to a final concentration of 292 mM NaCl (Thermo Fisher, cat. No. AM9759), 42 mM MgCl_2_ (Sigma-Aldrich, cat. No. M1028), 20 mM Tris (Sigma-Aldrich, cat. No. P7949), and CaCl_2_ (Sigma-Aldrich, cat. No. 10043-52-4). TST buffer was prepared containing 1zST buffer, 0.3% Tween-20 (Sigma-Aldrich, cat. No. P7949), 0.5% BSA (Miltenyi Biotec, cat. No. 130-091-376), and 1U/ml RNase inhibitor (Sigma-Aldrich, cat. No. 3335402001). 1X Lysis buffer was prepared on ice containing 1 % BSA (Miltenyi Biotec, cat. No. 130-091-376), 10 mM Tris HCL (pH 7.4, Sigma-Aldrich, cat. No. T2194), 0.01% Tween -20 (Sigma-Aldrich, cat. No. P7949), 0.01% Nonidet P40 Substitute (Sigma-Aldrich, cat. No. 74385), 3 mM MgCl_2_ (Sigma-Aldrich, cat. No. M1028), 10 mM NaCl (Thermo Fisher, cat. No. AM9759), 0.001% digitonin (Thermo Fisher, cat. No. BN2006), 1 mM DTT (Sigma-Aldrich, cat. No. 646563), 1 U/μl. RNase inhibitor (Sigma-Aldrich, cat. No. 3335402001). Lysis Dilution buffer contained 1% BSA, 10 mM Tris-HC (pH7.4), 3mM MgCl2, 10 mMNaCl, 1mM DTT and RNase inhibitor. Wash buffer was prepared to a final concentration of 1% BSA, 10 mM Tris-HCL, 0.1% Tween-20, 3 mM MgCl_2_, 10 mM NaCl, 1 mM DTT, 1 U/μl RNase inhibitor. A 25-50 mg frozen tumor sample was transferred to a 1.5 mL tube (Eppendorf, cat no.022431021) on dry ice before processing. Tissue was chopped in TST buffer using Noyes Spring scissors (Fisher Scientitic cat No. N9019530) for 10 minutes on ice. The homogenized solution was filtered through a 30 µm MACS SmartStrainer (Miltenyi Biotec, cat. No. 130-098-458) into a 15-mL tube. An additional 1 mL of TST was used to rinse the eppendorf and 3 mL of ST buffer to rinse the filter. The 5 ml homogenized volume was spun for 5 min at 500 g at 4°C. The supernatant was discarded, and the pellet was resuspended in 0.1X lysis buffer for 2 minutes on ice and then immediately rinsed with 1 ml wash buffer and spun for 5 minutes at 500 g at 4^0^C. The supernant was discarded and the nuclei pellet was resuspended in buffer, containing 1x nuclei buffer (20X nuclei buffer 10x Genomics, PN200207), 1 mM DTT, and 1U/μl RNase inhibitor. Final nuclei resuspension concentration was aimed 3,230-8,060 nuclei/μl with the target recovery maximum aimed at 10,000 nuclei. Cell count was performed using AO/PI staining and quantified using LUNA-FL™ Dual Fluorescence Cell Counter.

### Single nucleus RNA-sequencing

Transposition, GEM Generation and GEX libraires were prepared according the 10x Genomics Chromium Next GEM Single Cell Multiome ATAC + Gene Expression Guide CG000338. Approximately 8,000-10,000 nuclei per sample were loaded into each channel of a Chromium single-cell J chip (10x Genomics) and partitioned into droplets using 10x Chromium Controller to form emulsions, after which nuclear lysis, barcoded reverse transcription of mRNA, generation and amplification of complementary DNA, and 5’ adapter and sample index attachment were performed according to standard manufacturer’s protocol. All single-cell libraries were sequenced using an Illumina NextSeq 2000 sequencer.

### snRNA-seq data processing

CellRanger ARCv2.0 was used to demultiplex FASTQ reads, align them to the human reference genome (GRCh38) and extract the unique molecular identifiers (UMI) and nuclei barcodes. The output of this pipeline is a gene-barcode data matrix for each sample with the number of UMIs aligned to each gene. Cellbender ^65^ was used to remove ambient RNA and other technical artifacts from the count matrices. The parameters “expected-cells” and “total-droplets-included” were chosen for each dataset based on the total UMI per cell versus cell barcode curve in accordance with CellBender documentation. The false positive rate parameter was set to 0.01 by default. The false positive rate parameter “fpr” was set to 0.01.

Scrublet package (Python, version 0.2.3)^66^ was used for detection and removal of multiplet artifacts due to droplet encapsulation of more than one cell. An expected doublet rate was 0.05 and manually selected doublet score thresholds used to distinguish singlets from doublets in the bimodal score distribution graph. A singlet-only and ambient RNA-free count matrix was created for downstream analysis. Further quality control, dimensionality reduction, unsupervised clustering, and differential gene expression analysis were performed using Scanpy toolkit (version 1.9.2).

### Dimensionality reduction, clustering and cell-type annotation

Cells with fewer than 200 genes or greater than 5% counts representing mitochondrial genes were removed from each sample. Genes detected in fewer than three cells across all samples were also excluded. Raw counts data from responders and non-responder specimens were then normalized, scaled using default log-normalization parameters from Scanpy and aggregated into a single dataset. Linear dimensional reduction was performed on the integrated and normalized count matrix by performing principal components analysis (PCA) on the 5000 most highly variable genes in our dataset. The first 50 principal components were used for Leiden clustering of cells with a resolution parameter of 0.5. Uniform manifold approximation and projection (UMAP) was then performed using the same principal components with 100 nearest neighbors to visualize preliminary cell clusters from the nonintegrated dataset in two dimensions. To classify cells into broad cell types, differential expression analysis was performed comparing cells in each Leiden cluster to all other clusters using a two-sided Wilcoxon-rank sum test with Bonferroni FDR correction (q<0.01 considered significant). All samples were merged and integrated into a single data set using Harmony ^22^ to reduce technical variation and preserve biological diversity. The resulting integrated dataset included 59,006 nuclei and 12,169 detected genes across the 14 samples of the cohort.

### Identification of malignant cells

Cancer cells were identified using transcriptome inferred CNV profiles and cluster-level marker genes from the same tumor. Each sample underwent Leiden clustering and view map projection to identify *PTPRC^+^* (CD45^+^) and *PTPRC^−^* clusters. We then performed inferCNV^67^ within each sample using *PTPRC^+^* as a reference group and *PTPRC^−^* as observation groups. Malignant clusters were identified based on established copy number features of MIBC^11,12^ and combined to identify “tumor” cells within our cohort.

### Differential expression and gene set enrichment analysis

Differential expression analysis comparing cells from response groups (persistent tumor vs tumor scar) was performed using a two-sided Wilcoxon rank -sum test with Bonferroni FDR correction. We used fgsea v1.32.2^62^ for gene set enrichment analysis (GSEA). The log_2_(fold change) values for each gene were as ranks for pre-ranked for GSEA that was performed using as input all Hallmark gene sets from version 6.2 of the MSigDB^63^ or select gene sets curated from literature^48^. Gene sets with p < 0.05 and FDR < 0.25 were considered significant. Single-cell signature scoring was performed using VISION R package (version 3.0.0)^68^. Previously described macrophage, monocytes and DC markers described in the pan-cancer tumor infiltrating myeloid cell atlas^23^ were used to score bladder myeloid compartment (Fig. 3).

### Consensus non-negative matrix factorization

We utilized consensus non-negative matrix factorization (cNMF) to infer gene expression from programs from single-cell RNAseq data from bladder tumors ^28^. Gene expression matrices from both responders and non-responder samples were used as input. We performed 20 iteration per cell-type category and computed a set of consensus programs by aggregating results from all 20 runs and computed a stability and reconstruction error. Each program was annotated utilizing known described markers in the literature.

### Survival analysis of bulk RNA-seq data

Bulk RNA-seq data from two MIBC cohorts (GREEK, n=62; HM, n=55) treated with platinum chemotherapy (cisplatin and carboplatin) with OS annotated was obtained^10,34^. Patients who received carboplatin were excluded from this analysis, yielding a total of 66 cisplatin treated patients. Clinical and bulk RNA-seq data from TCGA was publicly available and used as a comparison group. To score malignant cNMF programs in each tumor, we summed the expression of the top 200 genes (Suppl Table 2) for each program and z-scored normalized the expression scores within GREEK/HM cohorts. Age, sex, and tumor stage were available for all patients. We used cNMF gene expression programs (usage 1, usage 2 etc) as aggregated programs to avoid model overfitting. Multivariable cox regression analysis was performed for OS in R 4.4.1.

### Receptor-ligand interaction inference

CellPhoneDB^53^ was used to infer receptor-ligand interactions. The algorithm was run on log-normalized expression values for malignant cells and tumor-associated macrophages.

### Tissue processing for spatial transcriptomics

RNA was extracted from the sample blocks to assess its integrity by running a DV200 assay for FFPE samples or RIN (RNA integrity number) measurement for fresh-frozen samples. For 10x Visium spatial transcriptomic assays, 5 µm FFPE or 10 µm fresh frozen sections were loaded on a Fisherbrand Superfrost Plus slide based on 10x Genomics Protocol CG000518. Subsequently, the slides were deparaffinized and stained for H&E. Tissue sections were scanned by a Nikon Eclipse Ti2 microscope for histological record then transferred to decrosslinking. Visium Human Transcriptome Probe Set was used and hybridized on a BioRad C1000 thermal cycler. RNA was then digested, probe ligated, released and extended. qPCR was used to quantify the eluted probes before running the sample index PCR and its cleanup. Libraries QC was performed on a Qubit 4 and Agilent Bioanalyzer 5200 and sequenced on the Illumina NovaSeq 6000 system or a Singular Genomics G4 with paired end reads according to the manufacturers guidelines.

### Spatial transcriptomics data processing

All persistent tumor speciments profiled for spatial transcriptomics (10x Visium) had matching snRNA-seq and spatial data was deconvoluted based on single nucleus cell-type signatures. We mapped the expression of each cell type to individual spots to label cell-types within the tissue sections. High resolution tiff images of the samples were annotated by a board-certified pathologist using the Samui browser. We used Cell2Location^69^ for single-cell reference-based spot deconvolution. We quantify a total of 11169 Visum spots each of which contain a median of 2718 unique genes after QC and decontamination with SpotClean. Subsequently, the Samui.json polygons with the pathologist the annotations (Suppl Fig. 4A) were used to subset spots using sf (1.0.14) and intersect that with the proportions, visualizing them with pheatmap (1.0.12). Visualizations for both single-cell and Visium data were created using ShinyCell (2.1.0) with the 10x Genomics VISIUM. Spatial domains to identify the co-existence and determine the proximity between SPP1/PARP14-Hi TAM and cancer urothelial cells was defined by SpaGCN^54^. Unsupervised iterative clustering (100 iterations per sample) that integrated spatially variable genes (SVGs), spatial location and histologic features defined a spatial domain overlaying the tissue area enriched in TAMs (Suppl. Fig 4). LIANA+^55^ was used for spatially inferred cell-cell interactions.

### Proximity ligation assay

Formalin-fixed paraffin-embedded (FFPE) tumor sections were deparaffinized in xylene and rehydrated through a graded ethanol series. Antigen retrieval was performed by boiling sections in citrate buffer (pH 6) for 20 minutes, followed by PBS washes and permeabilization with 0.5% Triton X-100 for 10 minutes. Sections were then blocked with 3% BSA in 0.1% Triton X-100 for 1 hour at room temperature. The sections were incubated with anti CD44 (1:100) and/or anti-SPP1 (1:100) overnight at 4°C. Subsequent steps were performed according to the manufacturer’s protocol using the Duolink® In Situ Detection Kit (DUO92008). Briefly, sections were washed twice with Wash Buffer A (5 min each), then incubated with PLA probes (anti-mouse PLUS and anti-rabbit MINUS; 1:5 dilution in blocking buffer) for 1 hour at 37°C. After washing, ligation was carried out using ligation stock (1:5) and ligase (1:40) in high-purity water for 30 minutes at 37°C. Sections were washed twice in Wash Buffer A (2 min each), followed by amplification with amplification stock (1:5) and rolling circle polymerase (1:80) for 1 hour 40 minutes at 37°C. Following amplification, sections were washed twice in Wash Buffer B (10 min each) and once in 0.01× Wash Buffer B. Slides were mounted with ProLong™ Diamond Antifade Mountant containing DAPI, and images were acquired using an Olympus FV3000R resonant scanning confocal microscope with a 63× objective.

### Multiplex immunofluorescence

Following sectioning, we incubate tissues at 60° C for 2 hours and followed a standard deparaffinization protocol. Antigen retrieval was done for 15 minutes at 95° C in a pressure cooker (BIO SB, Santa Barbara, CA, USA) in EDTA buffer (pH 8), then we quenched autofluorescence in bleaching solution (50mL PBS, 0.8 mL 1M NaOH and 4.5 mL H2O2) under LED light panels for 45 minutes. Samples were blocked for 1 hour under room temperature and incubated in fluorophore-conjugated antibodies for 16-18 hours at 4° C. Following antibody incubation, we rinsed tissue then fixed in 1.6% PFA to fix antibodies in place. We sealed overnight in ProLong Diamond Antifade Mountant with DAPI (Life technologies, Carlsbad, CA, USA). To quantify cell populations and proximity, we imaged three representative regions of each tissue section representing on a Zeiss Axio Observer 7 microscope (Zeiss, Oberkochen, Germany) with 20X objective. Regions were selected blind to sample group. Regions were stitched and processed with Zeiss Zen blue software prior to analysis.

### Macrophage cell culture and PARP14 assay

THP-1 and RAW264.7 cells were purchased from ATCC and cultured in DMEM medium (Gibco; 11965092) containing 10% heat-inactivated fetal bovine serum (Corning) and Penicillin/Streptomycin (Gibco; 15140122). THP-1 cells were differentiated into macrophages with 10ng/mL phorbol 12-myristate 13-acetate (PMA) for 72hr. Then, cells were treated with 0.1 μM of cisplatin for 72hrs, 0.1 μM of PARP14i (RBN012759; MedChemExpress; HY-136979) every 24hrs and combination cisplatin plus PARP14i. At the end of treatment RNA was extracted from lysed cells for RNA-sequencing (see below).

For RAW264.7 cells PARP14i (RBN012759; MedChemExpress; HY-136979) *in vitro* experiments, RAW264.7 cells were treated with 0.1µM PARP14i for 24 hours *in vitro*. Where indicated, RAW264.7 cells were treated with 20ng/mL recombinant mouse IFNγ (Biolegend; 575302) for the final 18 hours of PARP14i treatment *in vitro*. For Cisplatin *in vitro* experiments, RAW264.7 cells were treated with cisplatin for 72 hours *in vitro* using the indicated concentrations. RAW264.7 cells were assessed for PARP14 protein levels using flow cytometry. RAW264.7 cells were harvested from culture plates using TrypLE Express (Gibco; 12605036) and blocked for non-specific antibody binding using True-Stain Monocyte Blocker (Biolegend; 426101) and TruStain FcX Block (Biolegend; 101319) in FACS buffer (PBS + 1% FBS) for 10 minutes at room temperature. Cells were fixed and permeabilized using the Cytofix/Cytoperm Kit (BD; 554714) according to manufacturer’s instructions and stained for PARP14 (Santa Cruz; sc-377150) for 1 hour at 4C. Cells were acquired on an Aurora Spectral Flow Cytometer (Cytek) and analyzed using OMIQ (Dotmatics).

### Mouse experiments, treatments and tumor harvest

All animal experiments were performed in accordance with Dana-Farber Cancer Institute (DFCI) IACUC guidelines at the DFCI Longwood Center Animal Resource Facility under an approved protocol. The BBN963 bladder cancer cell line was a kind gift from William Kim (University of North Carolina). The PARP14 inhibitor RBN012759 was purchased from MedChemExpress (MCE) and dissolved in 0.5% methylcellulose (MC) plus 0.2% Tween 80. Cisplatin was purchased from the DFCI Research Pharmacy. Seven-week-old C57BL6/J mice (Jackson Labs) were subcutaneously implanted with 4.5 million BBN963 bladder cancer cells suspended in 75 ul PBS mixed 1:1 with Matrigel (Corning, 345230). Tumor volumes were monitored with digital calipers. When the tumor volume reached approximately 100 mm³, mice were randomized to one of four groups (9 mice per group) and treated with: (1) vehicle control (150 uL PBS injected intraperitoneally (i.p.) three times per week for 3 weeks and 250 µL of solvent (0.5% MC plus 0.2 tween 80) delivered via oral gavage (OG) twice per day for 3 weeks); (2) cisplatin alone (cisplatin 1 mg/kg in 150 uL injected i.p. three times per week for 3 weeks and 250 uL solvent (0.5% MC plus 0.2 tween 80) delivered via OG twice per day for 3 weeks); (3) PARP14 inhibitor alone (PARP14 inhibitor 500 mg/kg delivered via OG twice per day for 3 weeks and 150 uL PBS injected i.p., three time per week); and (4) cisplatin plus PARP14 inhibitor (cisplatin 1 mg/kg in 150 uL injected i.p. three times per week for 3 weeks and PARP14 inhibitor 500 mg/kg delivered via OG twice per day for 3 weeks). Tumor volumes were measured with digital calipers three times per week and mouse weights were measured twice per week during treatment and for an additional 11 days. At the end of the experiment, tumors were fixed in formalin and embedded in paraffin. Half of each tumor was sectioned for H&E staining and the other half was processed for bulk RNA sequencing.

### Bulk RNA-seq data

THP-1 cells and mouse BBN tumors were sent to Novogene (Durham, NC) for RNA extraction and sequencing (20 million reads per sample). The Nextflow RNA-seq workflow (nf-core/rnaseq; version 3.9) ^70^ was used to perform quality control, and STAR aligner and Salmon were used for alignment and reads quantification^71^. Counts were normalized using the median of ratios method, differential expression analysis was performed with DESeq2^72^ and GSEA using MiSgDB^62^.

## Data and code availability

Code and files used to generate all figures can be found at https://github.com/CarvalhoFilipeL/MIBC_tumor_myeloid and bulk RNA-seq cohorts used for single-nucleus RNA-seq tumor-intrinsic gene programs (Fig. 2D&E) in https://github.com/CarvalhoFilipeL/bladder_cancer_B_cell_sigs. Raw single-nucleus RNA-seq, spatial transcriptomic and bulk RNA-seq data will be deposited to dbGaP at the time of publication.

## Acknowledgments

We thank the patients who participated in this study and their families. This work was supported by the National Cancer Institute (R01 CA279221 (E.M.V.A., K.W.M.), P30 CA006516 (F.L.F.C.), K08 CA282969 (F.L.F.C.)), Parker Institute for Cancer Immunotherapy (E.M.V.A.), Bladder Cancer Advocacy Network (F.L.F.C.) and Karin Grunebaum Cancer Research Foundation (F.L.F.C.).

